# A novel High-Throughput Ligase-Independent Mapping method to detect Viral Integration Sites

**DOI:** 10.64898/2026.06.11.731714

**Authors:** Manisha Kabi, Ina Anreiter, Guillaume J. Filion

## Abstract

Integrated DNA elements are central to virology, functional genomics, and gene therapy, but current insertion-site mapping methods often rely on restriction digestion, ligation, or complex sequencing workflows that introduce bias and limit recovery. Here, we present Terminal Mapping, a high-throughput method that identifies host–insert junctions without restriction enzymes or DNA ligation. The workflow combines linear amplification from a known terminal sequence, enrichment of single-stranded products, terminal transferase-mediated poly-A tailing, and PCR amplification for Illumina or Oxford Nanopore sequencing. Applied to Jurkat T cells transduced with HIV-1- and SIVmac251-derived vectors, Terminal Mapping recovered more HIV-1 insertion sites than inverse PCR, reproduced known integration biases, and showed improved robustness with long-read sequencing. It revealed shared but quantitatively distinct HIV-1 and SIVmac251 hotspots, as well as substantial differences between Jurkat cell sources. Comparison of 5′ and 3′ LTR-derived reads also provided an internal control for unintegrated viral DNA. Terminal Mapping therefore offers a rapid and flexible platform for profiling integrated genetic elements across vectors and cellular contexts.

## INTRODUCTION

Integrated DNA elements are central to both genome biology and biotechnology. Retroviral vectors, transposons, and naturally occurring integrated viruses can modify the genome of their host through various integration mechanisms (Madhani, 2013). The range of phenotypic effects is remarkably wide, including viral replication, acquisition of pathogenicity or antibiotics resistance, deleterious mutations, or no effect whatsoever, but the genetic effect always amounts to the addition of a DNA sequence into a genome that is much larger in comparison (Hickey, 1982).

Integration is rarely neutral with respect to genome organization. Lentiviral systems, for instance, preferentially integrate into transcriptionally active, gene-rich regions and tend to avoid lamina-associated domains. Meanwhile, other integrating elements may show distinct preferences for promoters, enhancers, open chromatin, repetitive regions, or specific nuclear compartments. These biases matter for two reasons: First, they provide mechanistic insight into how integrating enzymes, host cofactors, nuclear architecture, and chromatin state influence insertion-site selection. Second, they determine the practical risks and limitations of integrating technologies. Insertions near oncogenes, tumor suppressor genes or regulatory elements can affect cell behavior, complicate interpretation of experimental models, and raise safety concerns in therapeutic applications (Bulcha et al., 2021).

For this reason, insertion-site mapping has become a core technology with applications in multiple fields. In gene therapy, it provides a method to assess vector safety by identifying integration hotspots near potentially hazardous loci and by monitoring clonal expansion. In genomics, it allows transposon or vector insertions to be connected to phenotypic outcomes. In virology, it provides a way to study persistent viral DNA, latent reservoirs, and the cellular history of infected clones. Finally, in single-cell studies, insertion sites can also serve as molecular barcodes, linking genomic events to cell identity or lineage structure. A broadly useful mapping method should therefore be sensitive, scalable, quantitative, compatible with different integrating elements, and minimally biased by local sequence composition.

Over the past two decades, integration-mapping technologies have evolved from low-throughput, locus-specific assays to genome-wide sequencing-based approaches. Early strategies such as inverse PCR relied on restriction enzyme digestion, circularization by ligation, and amplification across the unknown host–insert junction (Ochman et al., 1988; Triglia et al., 1988). These methods established a powerful conceptual framework: a known sequence within the integrated element can be used as an anchor to recover the unknown flanking genomic DNA. However, their practical performance is constrained by several features of the chemistry. Restriction enzyme dependence limits recovery to fragments containing suitably positioned cleavage sites, creating uneven coverage across the genome and reducing sensitivity in restriction-poor or repetitive regions. Ligation-dependent steps further reduce efficiency because the T4 DNA ligase suffers a catastrophic slow-down when the reaction reaches 30% completion. This imposes hard limits on the recovery when insertions are rare or even unique.

Later methods, including ligation-mediated PCR and linear amplification-mediated PCR, improved sensitivity and enabled broader use in retroviral and gene-therapy studies (Mueller & Wold, 1989; Schmidt et al., 2007). These approaches remain important, but many still depend on restriction digestion, adaptor ligation, nested amplification, or other steps that can introduce molecular bottlenecks. More recent short-read and long-read sequencing strategies have extended insertion-site mapping to higher throughput, richer genomic context, and in some cases single-cell resolution. Nevertheless, cost, complexity, throughput, and bias remain practical barriers, particularly for laboratories that need to profile many samples, compare vectors or cell types, or recover integrations across diverse genomic compartments.

Here, we present Terminal Mapping, a high-throughput, restriction enzyme-independent and ligase-independent strategy for mapping integrated DNA elements. The method is based on linear amplification from a known terminal sequence followed by A-tailing with the terminal transferase. The name Terminal Mapping reflects the central role of terminal transferase in the workflow as a replacement for DNA ligation, and the fact that the method targets the terminal junction between an inserted element and the host genome.

We developed Terminal Mapping as a general biotechnology platform for insertion-site discovery. The approach is designed to map the genomic positions of integrated vectors, transposons, or viruses using either Illumina short-read sequencing or Oxford Nanopore long-read sequencing. By avoiding restriction digestion, the method reduces sequence-dependent recovery bias and improves access to genomic regions that are poorly represented by enzyme-based protocols. By eliminating ligation, it avoids a major inefficiency of classical inverse PCR and related workflows, improving practical recovery of insertion sites from complex samples.

To evaluate the method, we applied Terminal Mapping to integrated lentiviral constructs in cell models and compared insertion profiles generated from HIV-1-and SIVmac251-based vectors. These experiments identified vector-and context-dependent integration patterns, including recurrent hotspots in Jurkat T cells. They also revealed that HIV-1-derived vectors can produce distinct insertion profiles in Jurkat T cells obtained from different sources, highlighting how integration-site selection may vary substantially across closely related cellular contexts. These observations reinforce the need for mapping approaches that are sensitive enough to capture sample-specific insertion landscapes rather than relying on assumed vector-level averages.

The complete Terminal Mapping workflow, from DNA extraction through library preparation can be completed within approximately two days. The option to sequence the libraries with the Oxford Nanopore technology typically allows for a rapid evaluation of the result. This fast turnaround connects experimental work directly to genomic interpretation and makes the method suitable for iterative vector design, comparative benchmarking, safety assessment, and exploratory studies of integrated DNA biology. More broadly, Terminal Mapping provides a practical route for laboratories to profile the composition, distribution, and dynamics of inserted genetic elements across experimental systems.

## MATERIALS AND METHODS

### Cell culture

The human Jurkat T cell line was maintained in RPMI 1640 medium (Gibco, 11875-093) supplemented with 10% fetal bovine serum (FBS) (Corning, 35-077 CV), 1% GlutaMAX (Gibco, 35050-061), 100 U/ml penicillin-streptomycin (Gibco, 15140-122), and 1% MEM Non-essential Amino acid (Gibco, 11140-050). HEK 293T cell line was cultured in Dulbecco’s modified Eagle’s medium (DMEM) (Gibco, 11965-092) supplemented with 10% FBS. All cells were incubated at 37°C in the presence of a 5% CO_2_ atmosphere and passaged every three days with a 1:5 dilution. One batch of the Jurkat T cell line (Jurkat-a) was kindly provided by L. Di Croce (Center for Genomic Regulation, CRG, Barcelona). The other batch of Jurkat T cell line (Jurkat-b, known to be a Jurkat E6-1 clone) and HEK293T cell line were kindly provided by C. Guzzo (University of Toronto Scarborough, Toronto).

### Construction of HIV-1 and SIVmac251 mini vectors

All HIV-1 transductions were carried out in a single-round with a replication-deficient HIV-1 construct derived from plasmid pEV731 which was kindly provided by A. Jordan (IBMB, Barcelona). pEV731 is an HIV-based vector encoding the two-exon form of the HIV-1 tat gene and a GFP marker gene under control of the HIV-1 LTR. The sequence of HIV-1 with schematic structure “LTR-Tat-IRES-EGFP-Nef-LTR” was PCR-amplified from pEV731 and cloned into a 2.9kb high-copy number backbone pUC19 for propagation in *Escherichia coli*. GFP minimal SIVmac251 transfer vector, was received from F. Margottin-Goguet (Institut Cochin, Paris). The vector was constructed based on the previously described gaMES4 vector carrying a self-inactivating design (Mangeot et al., 2002) and a CMV-eGFP cassette downstream of SIVmac251 LTR (Goujon et al., 2006).

### Transfection and viral transduction in Jurkat T cells

HIV-1 and SIVmac251 pseudotyped viruses were generated using the standard calcium phosphate protocol (Jordan et al., 1996). Briefly, 2×10^6^ HEK 293T cells in 100 mm dishes were cotransfected with 6.5 μg pCMVΔR8.91 packaging plasmid, 3.5 μg pVSV-G and 10 μg HIV-1 viral vector or 6.5 μg pSIV3+Vpr+Vpx+ packaging plasmid, 3.5 μg pVSV-G and 10 μg SIVmac251 viral vector. Culture supernatants were harvested 48 hours post-transfection, filtered using a 0.45 μm filter, and stored in 1 mL aliquots at-80 °C prior to further use.

The relative infectious units (RIU) of the pseudotyped reporter viruses were quantified using 3×10^5^ Jurkat T cells in each well of a 12-well tissue culture plate. The cells were transduced with viral stocks serially diluted 2-fold (from 10x to 80x) in a total volume of 1 mL of complete RPMI 1640 medium containing 8 μg of Polybrene (MilliporeSigma, TR-1003). The plates were centrifuged at 1,000 g at 32 °C for 90 min. At 48 hours post transduction, the efficiency of transduction was analyzed by the proportion of GFP (+) cells using a flow cytometer (Becton Dickinson, LSRFortessa). Following this, the RIU / mL was calculated using dilutions resulting in < 20% infection to minimize multiple infection events per cell, using the formula: RIU / mL = [cell number x ratio of GFP (+) cells x dilution factor] / volume (mL). Subsequently, the multiplicity of infection (MOI) was defined as the ratio of the RIU to the number of host cells at the time of transduction experiment (MOI = RIU / cells). For all subsequent experiments, the volume of viral supernatant required to achieve a target MOI (*e.g.*, 0.1) was adjusted based on this calculated functional titer to ensure consistent infection efficiency across replicates.

### FACS sorting

Jurkat T cells were sorted two days post transduction with pseudotyped reporter viruses at an MOI of 0.1 with a cell sorter (Becton Dickinson, FACSAria III). Before sorting, cells were washed and resuspended in PBS (Gibco, 10010-023) containing 1% FBS and 5 μL of Hoechst dye (Thermo Scientific, 62249) at a concentration of 1 μg / mL. Results shown throughout the chapter correspond to representative data from experiments repeated at least twice.

### Nucleic Acid isolation

Genomic DNA from Jurkat T cell pools was extracted three weeks post-transduction with PureLink Genomic DNA Kit (Invitrogen, K1820-01). Genomic DNA was quantified on a NanoDrop (DeNovix Inc., DS-11FX) spectrophotometer.

### Library preparation of Terminal Mapping samples for High-Throughput sequencing

500 ng genomic DNA from transduced Jurkat T cell sub-clones were linear-amplified using either 0.02 μM CCL0042 or CCL0064 (annealing to the HIV-1 5′ LTR and to the SIVmac251 5′ LTR, respectively) and 1.1x Red Taq DNA Polymerase Master Mix (VWR Life science, 733-2544) in a 50 μL final volume. The cycling conditions were as follows: 95°C for 5 min; 95°C for 30 s, 55°C for 1 min and 72°C for 15 s (100 cycles); and 72°C for 5 min. 5 μL of the reaction volume was used to confirm linear amplification by PCR using either 0.2 μM CCL0025 and CCL0033 (annealing to the HIV-1 vector) or 0.2 μM CCL0067 and CCL0068 (annealing to the SIVmac251 vector) in 20 μL standard Taq DNA polymerase reaction mix (GeneBio Systems Inc., P3004). The cycling conditions were as follows: 95°C for 5 min; 95°C for 30 s, 60°C for 30 s and 72°C for 30 s (25 cycles); and 72°C for 5 min.

Following linear amplification, 25 μL single-stranded DNA product in the Taq DNA Polymerase Master Mix was mixed with either 0.4 μM CCL0034 (biotinylated primer annealing to CCL0042 for HIV-1) or 0.4 μM CCL0065 (biotinylated primer annealing to CCL0064 for SIVmac251) and hybridized by gradual cooling. The mixture was denatured at 95°C for 2 min, followed by slow cooling from 95°C to 15°C at a rate of 0.1°C/s. The temperature was held at 15°C for 10 min to ensure complete annealing. Biotinylated single-stranded DNA was then purified using streptavidin-coated magnetic beads (Cytiva, 30152103010350) according to the manufacturer’s instructions. Bead-bound DNA was washed thoroughly three times with 1x High salt wash buffer containing 10 mM Tris HCl pH 8.0, 1 M NaCl and 0.5 mM EDTA. The purified single-stranded DNA was eluted in 20 μL Elution Buffer (Qiagen, 19086) preheated to 95°C. The mixture was incubated at 95°C for 5 min to disrupt biotin-streptavidin interactions. The beads were then magnetically separated, and the supernatant containing the eluted DNA was collected.

To catalyze the addition of poly(A) tails to the 3′ ends of DNA fragments, terminal deoxynucleotidyl transferase (TdT) reaction was performed using the 20 μL eluted DNA, 5 μL 10x Terminal Transferase Reaction Buffer, 0.25 mM CoCl_2_, 1 μL 10 mM dATP, and 20 units TdT enzyme (New England Biolabs Inc., M0315) in a 50 μL final volume. The reaction mixture was incubated at 37°C for 30 min, followed by enzyme inactivation at 70°C for 10 min.

For exponential amplification by PCR, 25 μL of TdT reaction products were directly added in 50 μL standard Taq DNA polymerase reaction mix (GeneBio Systems Inc., P3004) with 0.2 μM forward primer CCL0045 (annealing to the poly(A) tail with Illumina R2 Seq Primer overhang) and 0.2 μM reverse primer either CCL0026 or CCL0066 (annealing to the HIV-1 5′ LTR and to the SIVmac251 5′ LTR, respectively; with Illumina R1 Seq Primer overhang). The cycling conditions were as follows: 95°C for 5 min; 95°C for 30 s, 53°C for 1 min and 72°C for 1 min (30 cycles); and 72°C for 5 min. For each condition, at least four tubes were pooled and purified using a silica column-based purification kit (Qiagen, 28104) according to the manufacturer’s instructions and eluted in 20 μL Elution Buffer. To add the Illumina primers, 4 μL of the purified product were diluted to 50 μL of standard Taq DNA polymerase reaction mix with 0.2 μM primer GAT024 (annealing to the Illumina PE1.0 primer) and an indexing primer GAT-int (annealing to the Illumina PE2.0 primer). The cycling conditions were as follows: 95°C for 5 min; 95°C for 30 s, 56°C for 1 min and 72°C for 1 min (6 cycles); and 72°C for 5 min. PCR products ran as a smear on agarose gel.

The specificity of the reaction was checked by ensuring that the smear was absent in reactions where the cells were not transduced (no viral DNA in the input, asserting that linear amplification does not proceed from off-target genomic sites) and where no linear amplification was performed (asserting that the biotinylated oligo does not anneal to off-target genomic sites).. All the primers used are described in Supplementary Tables 1 and 2.

*Oxford Nanopore sequencing.* Replicate libraries were pooled together if they needed to be sequenced in the same flow cell. The pooled libraries were adjusted to a 20 ng/μL final concentration. DNA Libraries were repaired using NEBNext FFPE DNA Repair Mix (NEB, M6630) and Ultra II End-prep (NEB, E7546) according to the manufacturer’s instructions. Sequencing adapters were attached to the DNA libraries using Ligation Sequencing Kit (SQK-LSK109) and MinION flow cells were primed using the Flow cell priming kit (EXP-FLP002) according to the manufacturer’s instructions. Libraries for mapping were sequenced with R9 flow cells (Oxford Nanopore Technologies, FLO-MIN106D) on a MinION Mk1B sequencer.

*Illumina sequencing.* Replicate libraries were pooled together if they needed to be sequenced in the same lane of the flow cell. The pooled libraries were adjusted to a 4 nM final concentration. Libraries for mapping were sequenced as a 75-bp single-read on a NextSeq sequencer with NextSeq 500 (400M clusters) high output kits at the Donnelly Sequencing Centre (https://thedonnellycentre.utoronto.ca/).

## Data analysis

*Sequence extraction from Nanopore reads.* Viral genomic segments were extracted from FASTQ files using a custom Python parser based on seeq for approximate DNA/RNA matching via Levenshtein distance (https://github.com/ezorita/seeq). Briefly, sequencing reads and their reverse complements were scanned for the anchor consisting of the adapter and the added poly-A tail with a maximum tolerance of 6 errors including indels (target sequence: AAAAAAAGATCGGAAGAGCACACGTCTGAACTCCAGTCAC). Identified reads were split around the anchor, using the prefix to find an insertion site in the genome, and recursively repeating the process above on the suffix.

The reason for recursively searching for insertion sites in the same read is that independent molecules are routinely ligated together to form concatemers during the adapter ligation step, so a single read may contain multiple amplicons. For each prefix, the specific viral motif from the LTR was identified with a maximum tolerance of 5 errors including indels (HIV-1 target sequence: CTTGTCTTCGTTGGGAGTGAATTAGCCCTTCCA, SIVmac251 target sequence: TCTATGTCTTCTTGCACTGTAATAAATCCCTTCCA). The sequence located between the viral LTR and the closest poly-A tract was isolated (20 As with a maximum tolerance of 3 errors including indels) and kept for further analyses. Given that the sequences of the 5’ and 3’ LTRs are identical, the chunks between a viral LTR and a poly-A tract are either the sequence of the host cell immediately upstream of the 5’ LTR (*i.e.*, it is the insertion site of the construct), or the sequence of the virus immediately upstream of the 3’ LTR.

*Sequence extraction from Illumina reads.* The ends of the LTRs were trimmed by seeq allowing 5 errors (HIV-1 target sequence: CTTGTCTTCGTTGGGAGTGAATTAGCCCTTCCA, or SIVmac251 target sequence: TCTATGTCTTCTTGCACTGTAATAAATCCCTTCCA), leaving 34 nucleotides (plus or minus errors) for mapping the read into the host genome.

*Mapping insertion sites in the host genome.* Processed Nanopore reads were mapped using bwa with options bwa mem ‘-t8-L0-x pacbiò in an index containing the human genome (hg38), the genome of phage PhiX174, and the sequence of the HIV and SIV constructs. The added reference sequences accelerate the mapping and allow us to perform some essential controls (see below). Following alignment, parsed SAM files were processed to identify high-confidence integration sites. Alignments containing hard clipping (secondary alignments) or exhibiting more than 5 bp of soft clipping at the 5’ end were discarded to ensure the sequencing read began immediately at the host-virus junction. Genomic coordinates were determined based on alignment orientation: for positive strand matches, the integration site was defined by the alignment start position, whereas for negative strand matches, it was defined by the alignment end (start position plus alignment length derived from the CIGAR string). To correct for sequencing or mapping imprecision, insertion sites on the same chromosome located within a 15 bp window were collapsed into a single one located at the mode of the cluster. Reads mapping to the internal controls (HIV, SIV, and PhiX174 genomes) were filtered out at this stage. Finally, the representative integration site for each cluster was assigned to the coordinate with the highest read count, and the number of independent capture events was estimated by quantifying unique poly-A termination sites (separated by 5 bp or more) associated with that insertion.

Processed Illumina reads were mapped using bwa with options ‘bwa mem-t8-L0‘ in the index described above. The mapping process was the same as for Nanopore reads, except for the last step because poly-A tracts could not be used to demultiplex independent linear amplification events. With only 34 nucleotides available for mapping, either the read did contain a poly-A tract but could not be mapped with high confidence for lack of remaining nucleotides, or the read did not contain a poly-A tract.

The scripts that were used to extract reads and map the viral insertions sites are publicly available on Github at https://github.com/gui11aume/terminal_mapping.

*Comparing reads from the 3’ and 5’ LTRs.* To compare the number of reads from the 5’ and 3’ LTRs, we filtered the mapped reads in the output of bwa (see Mapping insertion sites in the host genome). For this, we used‘samtools sort‘and‘samtools index‘to sort and index the SAM files produced by bwa, then we used‘samtools view‘with options‘-cq20‘ to count the number of reads with mapping quality 20 or above. We first discounted a common false positive at position 17402086 of chromosome 3 that was found in every experiment. The command ‘samtools view-cq20 [indexed-bam-file.bam] chr3:17402074-17402154‘ was used to compute a number x. We then counted the number y of reads with quality 20 or above in mini HIV or mini SIV (depending on the experiment), and finally we counted the number z of reads with quality 20 or above in the entire experiment. The proportion of reads from the 3’ LTR was reported as the ratio y / (z - x) converted to percent.

## RESULTS

### Principle of Terminal Mapping

To improve on previous insertion-mapping techniques, we developed Terminal Mapping, a high-throughput method to determine the integration sites of integrated viruses with high robustness. Briefly, the key features of Terminal Mapping are to initiate a linear amplification step from integrated viruses, purify the products by DNA hybridization, and use the terminal transferase to add a poly-A tail to the purified products. At that step, the products have known ends, with a viral LTR at one end, and a poly-A tail at the other, allowing them to be amplified by PCR. The poly-A tail replaces the ligation of an adapter with known DNA sequence, except that the efficiency of the terminal transferase is superior to that of DNA ligases. In addition, Terminal Mapping does not require any DNA fragmentation, which can waste large amounts of input material (when using sonication) or limit the range of mappable insertions (when using restriction enzymes).

The insertion patterns of HIV-1 have been extensively characterized in the clinic and in research (Lusic & Siliciano, 2017), but the related virus SIV has received substantially less attention in this regard. As a proof of concept for Terminal Mapping, we set out to compare the integration profiles of HIV-1 and SIVmac251 to determine whether SIVmac251 has integration hotspots and whether they are the same as those of HIV-1 (Crise et al., 2005; Ferris et al., 2019). To this end, we used a 4.6-kb minimal HIV-1 construct containing the *tat* transcriptional activator and a GFP-Nef-encoding fusion gene (**Fig. 1a, top panel**) and a 4.8-kb minimal SIVmac251 transfer vector containing GFP under the control of cytomegalovirus (CMV) promoter (**Fig. 1a, bottom panel**). We obtained infectious pseudotyped particles by cotransfection with a plasmid expressing VSV-G and transduced Jurkat T cells with a viral inoculum at a multiplicity of infection (MOI) of 0.1 (**Fig. 1b**). The efficiency of HIV-1 transduction was comparable to that of SIVmac251 (**Supplementary Fig. 1a**). HIV-1 and SIVmac251 transduced GFP (+) Jurkat T cells were sorted by flow cytometry and used to establish a pool of founder cells (**Supplementary Fig. 1b**). This cell pool was then expanded in culture so that each founder cell produced a clone with identical proviruses (**Fig. 1c**). The purpose of this approach was to ensure that every cell in the founder population is infected with at least one virus, while reducing the number of cells with more than one provirus.

**Figure 1.**
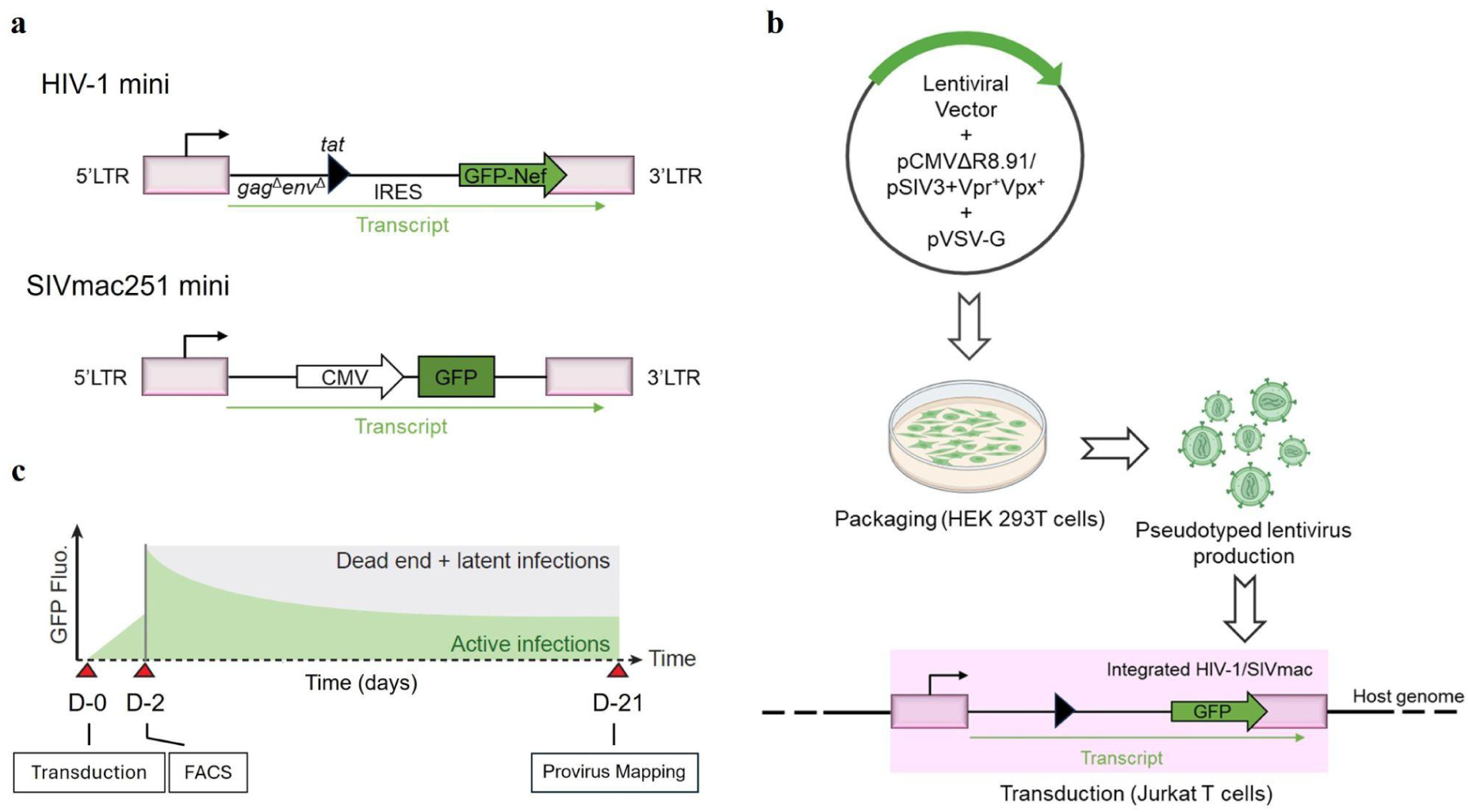
Structure of the parental minimal lentiviral vectors and transduction in Jurkat T cells. (a) Top: The minimal HIV-1 construct expresses Tat and GFP under the control of the HIV-1 LTR. IRES: internal ribosome entry site. Bottom: SIVmac251 minimal transfer vector bearing GFP under the control of CMV, cytomegalovirus promoter. (b) Infectious viral particles were prepared by co-transfection of HEK 293T cells with the minimal lentiviral vectors (HIV-1 and SIVmac251) and two additional plasmids expressing VSVG and the packaging proteins (pCMVΔR8.91/pSIV3+Vpr+Vpx+), respectively. Culture supernatants were harvested 48 hours post transfection. Jurkat T cells were transduced with the generated pseudotyped viruses. The region highlighted in pink shows the structure of an integrated provirus. (c) Experimental outline of HIV-1 or SIVmac251 transduction in Jurkat T cells. The cells were transduced with HIV-1 or SIVmac251 pseudotyped reporter viruses at an MOI of 0.1. GFP positive cells were FACS-sorted 2 days after transduction. The sorted cells were expanded for three weeks before mapping the viral insertions.

We then mapped the integrated proviruses by Terminal Mapping (Methods and **Fig. 2**). PCR products ran as a smear on agarose gel (**Supplementary Fig. 2**), representing the varying sizes of the amplicons containing the insertion sites. The smears failed to appear when the cells were not transduced or when no linear amplification was performed, showing that the products were not the result of non-specific amplification of background signal. Libraries were sequenced using Illumina and Nanopore technologies in order to compare mapping workflows with short and long reads, respectively.

**Figure 2.**
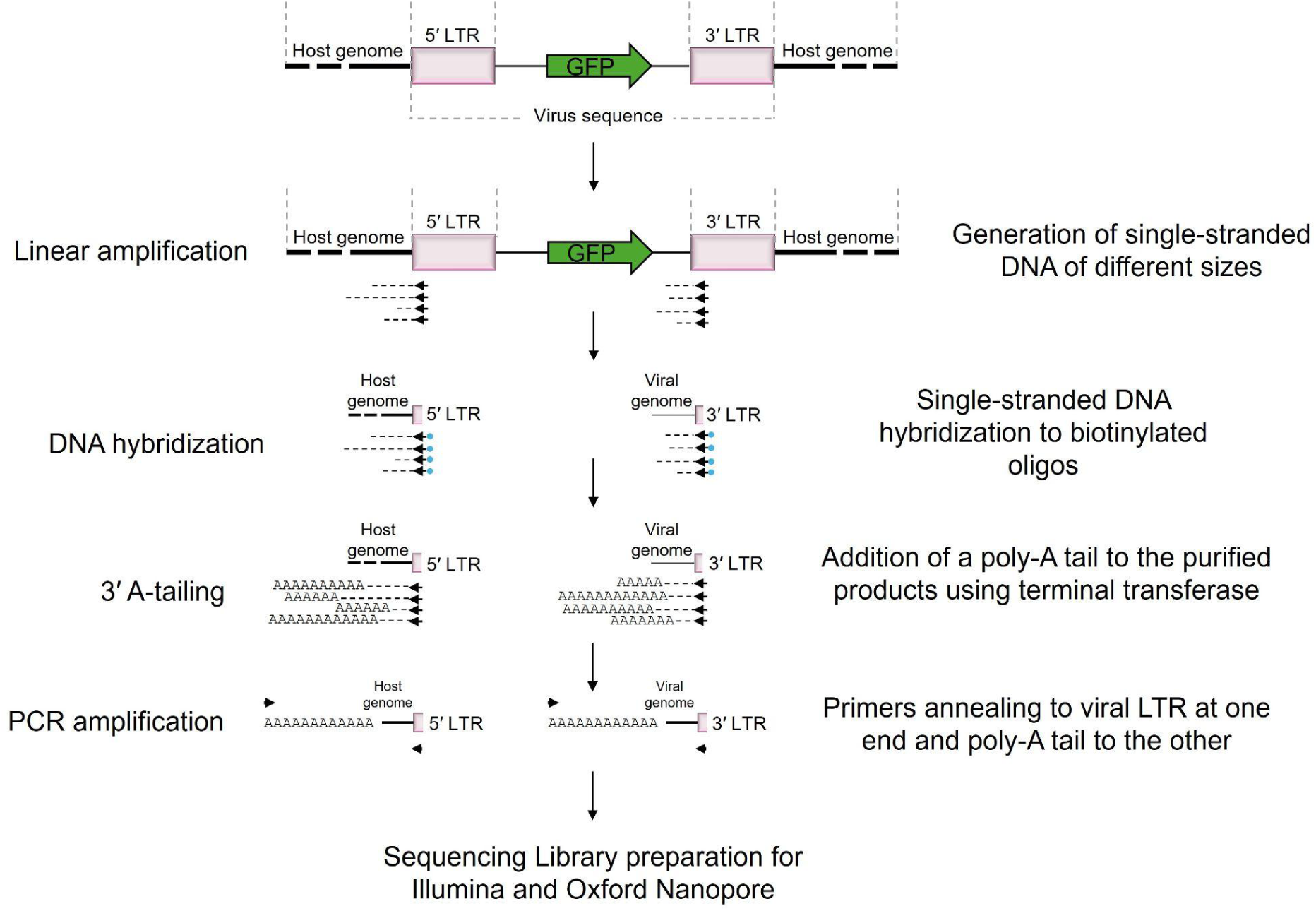
Schematic outline of Terminal Mapping. The method starts with an initial linear amplification of vector genome junctions using a primer hybridizing close to the end of the known DNA sequence: long terminal repeat (LTR) of a lentiviral vector. Preamplification results in ssDNA of different sizes. Amplified ssDNA targets are annealed to biotinylated oligos, captured on magnetic particles and purified. Terminal transferase (TdT) catalyzes the addition of dATPs to the 3’ hydroxyl terminus of the linear amplified DNA molecules generating a poly A-tail. The end step is exponential amplification by PCR with vector-specific primers containing sequencing adaptors.

### Terminal mapping recovers known insertion profiles

HIV-1 and SIV have a characteristic nonrandom integration pattern preferentially targeting active genes and gene-rich chromosomes in lymphoid cell lines (Schröder et al., 2002; Crise et al., 2005). Here, we first aimed to recapitulate the established features of HIV-1 integration that we previously characterized in Jurkat T cell line-a using inverse PCR (Chen et al., 2017). To enable a direct comparison with our earlier work, we first analyzed a biological sample generated at the same time as the inverse PCR dataset. Only one replicate of this sample was available, but it was important to compare the methods based on the same biological material for reasons that will be detailed below.

Terminal mapping recovered substantially more integration sites than inverse PCR under comparable experimental conditions. Inverse PCR identified 2,472 integration sites across two independent transductions, whereas terminal mapping identified 11,441 integration sites from a single transduction experiment. Overall, the recovery of Terminal mapping was approximately an order of magnitude above that of inverse PCR, suggesting that the various optimizations of Terminal mapping improve the overall efficiency of the workflow.

A genome-wide analysis of HIV-1 integration sites revealed broadly similar chromosomal insertion biases between the two methods. The preferential integration of HIV-1 into chromosomes 16, 17, and 19 has been well documented previously (Bushman et al., 2005) and was likewise observed here. Nevertheless, the chromosomal integration profiles differed significantly between methods (chi-square test with 23 degrees of freedom, P = 0.03), indicating that the two approaches do not generate strictly identical outputs. Despite this statistical significance, the relative differences in chromosomal insertion frequencies were modest, on the order of ∼10% (**Fig. 3a**), even for datasets exceeding 10,000 integration events. These results suggest that inverse PCR and terminal mapping each introduce method-specific biases, while preserving broadly comparable large-scale integration patterns.

**Figure 3.**
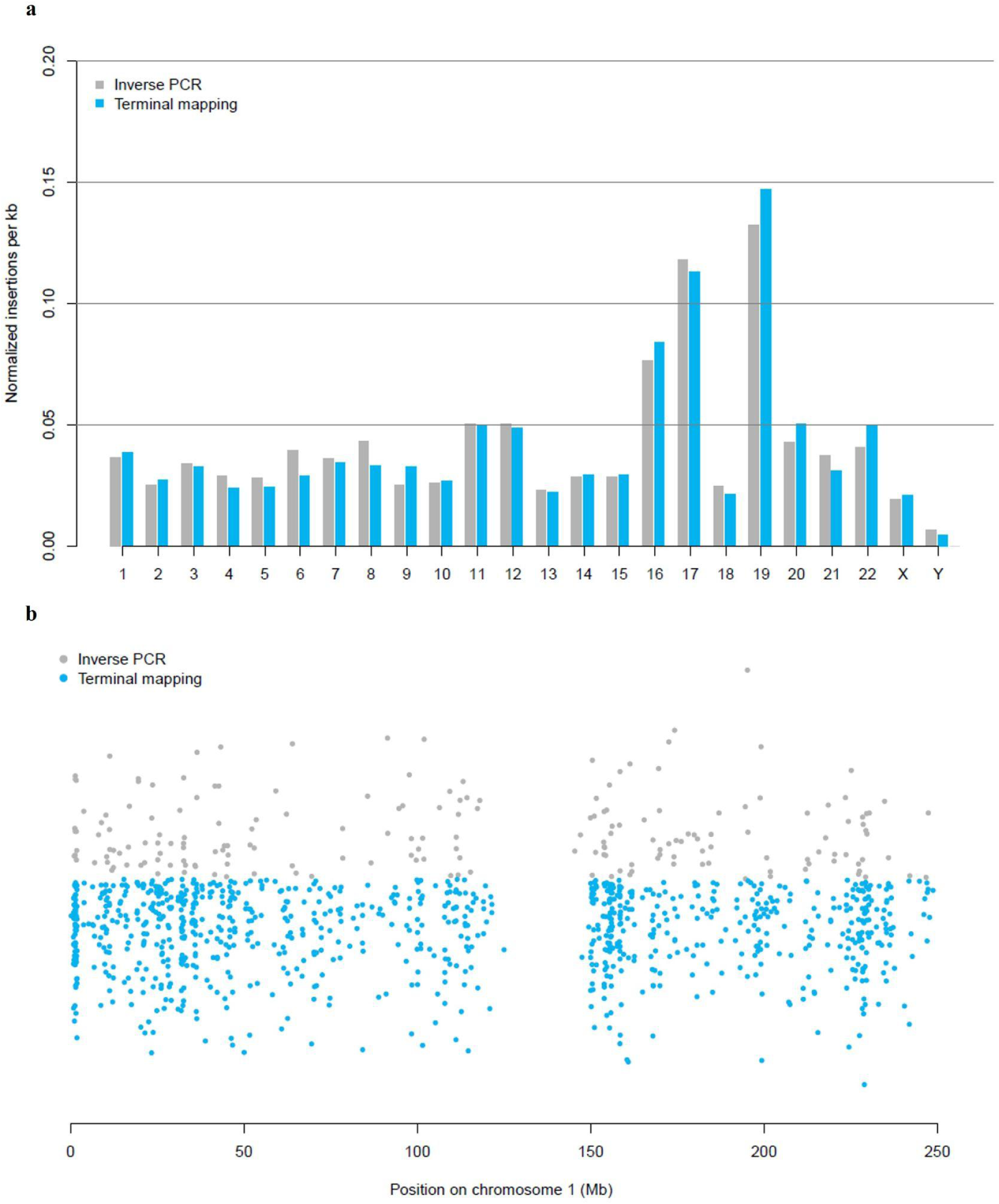
Inverse PCR vs Terminal Mapping. (a) Bar graph showing the number of mapped insertions per kilobase for the HIV-1 construct in Jurkat T cells (clone Jurkat-a), mapped with inverse PCR (Chen et al., 2017) or with Terminal Mapping (this study). The profiles and the insertion biases at the genome-wide scale are similar but statistically distinguishable (chi-square test with 23 degrees of freedom, χ =37.44, P = 0.03). (b) In this representation, each dot is an integrated provirus and its x-coordinate is the reported insertion site. The y-coordinate is a random jitter allowing nearby proviruses to be distinguished. In this representation integration hotspots appear as concentrations of dots on a near-vertical line. The insertion sites obtained by inverse PCR (in gray) point upward, while those obtained by Terminal mapping (in blue) point downward. There are more insertion sites with Terminal mapping than with inverse PCR, but the overall shape and the position of the hot spots are the same for both techniques.

To compare the insertion profiles at a finer scale, we plotted every insertion on chromosome 1 (**Fig. 3b**). In spite of the downward skew (due to the higher recovery of Terminal mapping) it is possible to identify the alignment of hotspots, showing that the profiles share the same overall features at chromosome scale. Several regions are not covered by any inverse PCR insertion site, but show clear coverage with Terminal mapping, albeit with lower density. This suggests that the increased coverage of Terminal mapping gives a higher resolution of the insertion profile.

### Terminal mapping leverages long-read sequencing

In the light of the results above, we were surprised to find large differences in the HIV-1 integration landscape when we compared two Jurkat T cell lines from different sources, Jurkat-a and Jurkat-b (Methods). For context, we noticed that our results in a new research environment were noticeably different from earlier results and we were able to pin down the source of the discrepancy to the Jurkat clone. Having no longer access to the clone Jurkat-a, we set out to use Terminal mapping to establish the baseline integration profile of Jurkat-b, using both HIV-1 and SIVmac251 constructs in order to gain insight into the distribution of integration hotspots. We also set out to test different sequencing technologies in order to establish the performance of Terminal mapping. We therefore carried out two independent HIV-1 and two independent SIVmac251 transductions in Jurkat-b, performed two Terminal mappings on each sample and sequenced the libraries on either the Illumina NextSeq or the Oxford Nanopore Technology MinION.

Short-read sequencing offers key advantages in terms of throughput and accuracy. High read output can improve recovery rates, particularly when sequencing libraries contain low proportions of target molecules (*e.g.*, because of contaminations). In addition, the high accuracy of short reads facilitates the reliable identification of viral sequences and facilitates the preprocessing of the reads. On the other hand, long reads have two important advantages for Terminal Mapping: First, they provide a larger window around the genomic context at the insertion site, making it easier to map the insertion site uniquely with software such as bwa or bowtie. Second, long reads (median 345bp, interquartile range 269–498 bp) typically contain the poly-A tail added by the Terminal Transferase (**Supplementary Fig. 3**), allowing the analyst to distinguish independent products of linear amplification. Specifically, linear amplification will terminate at different locations on every iteration, so the poly-A tail will also start at different locations. Capturing multiple independent events of linear amplification gives additional confidence that a virus was actually inserted at the given location. By contrast, most spurious events, such as contaminations or off-target PCR amplifications are based on a single initial molecule, in which case the poly-A tail, if any, is always at the same location in every read. In this regard, Terminal Mapping provides an additional indicator of the reliability of the results, provided the reads are long enough to capture the position of the poly-A tail.

With short reads, we recovered 5,179 + 4,676 integration sites for HIV-1 (transduction 1 plus transduction 2) and 13,319 + 5,673 integration sites for SIVmac251 (transduction 1 plus transduction 2). Using long reads, we recovered 7,228 + 6,618 integration sites in HIV-1 (transduction 1 plus transduction 2) and 19,467 + 9,353 integration sites in SIVmac251 (transduction 1 + transduction 2). In addition, we also used long-read technologies to sequence the Terminal mapping library from Jurkat-a (see above) and obtained 16,117 integration sites, compared to 11,441 with short-read technologies. In conclusion, using long-read sequencing increased the number of mapped integration sites for each single library.

However, the fact that there are more insertion sites does not imply that recovery is higher, as several insertion sites may be false positives originating from technical artefacts. To benchmark the sequencing methods on more informative measurements, we compared the replicates of Terminal mapping performed on the same input DNA, aiming to determine how the sequencing technology contributes to the reproducibility between technical replicates (*i.e.*, all experimental steps after harvesting the biological material). The number of insertion sites that were present in both replicates was higher with Nanopore than with Illumina (**Fig. 4a**). This was true for either HIV-1 or SIVmac251 constructs, suggesting that long reads delivered more high-quality insertion sites than short reads. For HIV-1 constructs, the proportion of insertion sites present in both replicates was similar between Illumina and Nanopore (32% vs 30%, respectively), but for SIVmac251 constructs, the proportion of insertion sites present in both replicates was 50% higher with Nanopore than with Illumina, was 21% vs 14%. Taken together, these results indicate that the higher number of insertion sites with long reads is not due to the accumulation of random noise. On the contrary, the proportion of high-quality insertion sites was higher in the case of SIVmac251.

**Figure 4.**
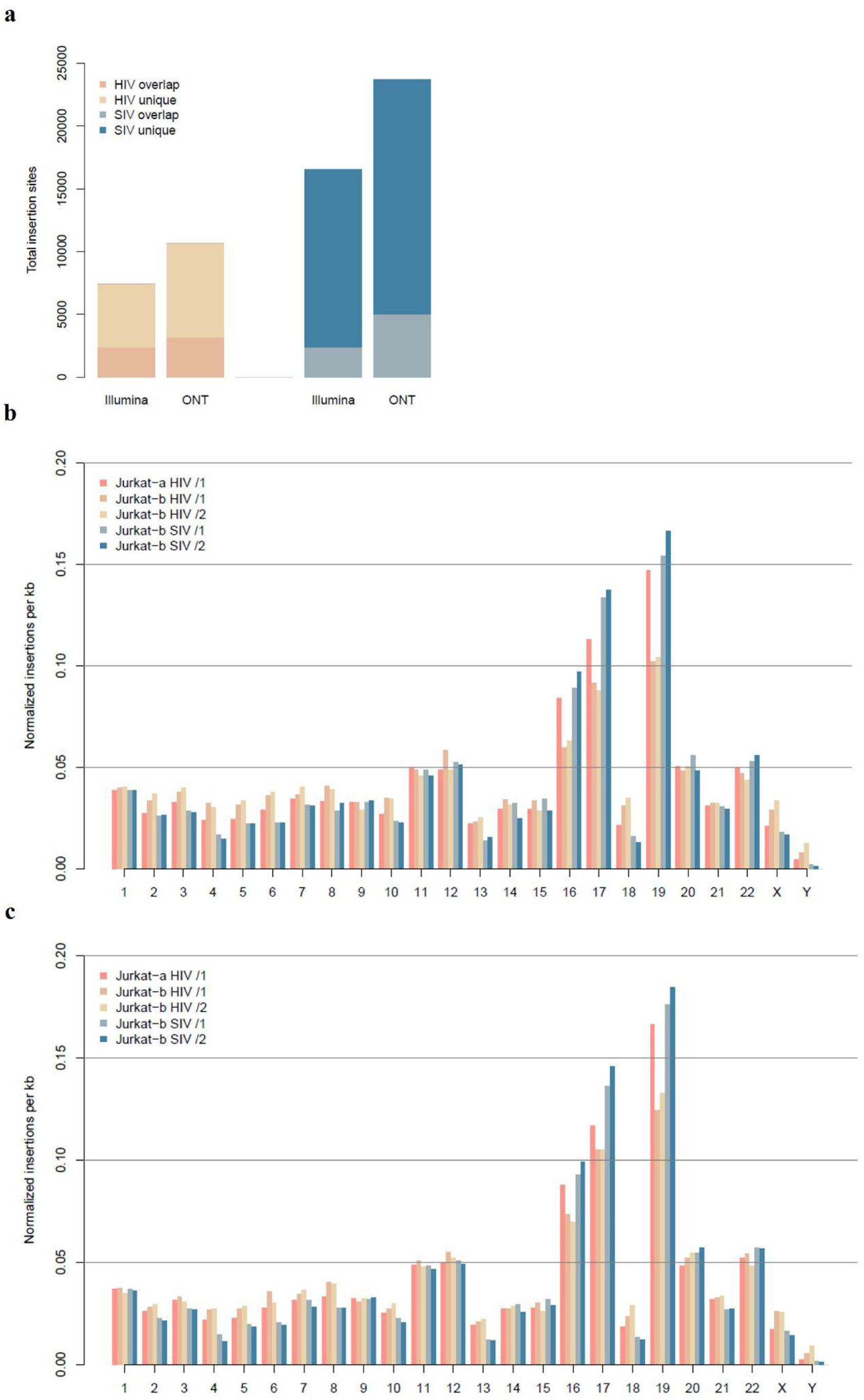
Integration landscape of HIV-1 and SIVmac251. (a) Stacked bar plot showing the number of insertions found in two replicates (overlap) or just one (unique) for HIV-1 (yellow) or SIVmac251 constructs (blue). The replicates were sequenced with either Illumina or Oxford Nanopore Technology (ONT). The number of insertions found in two replicates is higher with ONT. Libraries were sequenced using the Illumina sequencing technology (b) and Oxford Nanopore sequencing technology (c). For both (b) and (c), the number of mapped insertions per kilobase is represented as a bar graph for both HIV-1 and SIVmac251 transductions in Jurkat T cells. Jurkat-a and Jurkat-b are cell lines from different sources. The chromosomal distribution of integrated HIV-1 and SIVmac251 transductions showed a clear enrichment on chromosomes 16, 17 and 19. All three figures show one HIV-1 transduction in Jurkat-a, two independent HIV-1 transductions in Jurkat-b and two independent SIVmac251 transductions in Jurkat-b. The differences are statistically significant between Jurkat-a and Jurkat-b for the HIV-1 construct (chi-square test with 23 degrees of freedom, for (a) χ^2^=238.38, P < 0.001 and for (b) χ^2^=191.94, P < 0.001) and for the SIVmac251 construct (chi-square test with 23 degrees of freedom, for (a) χ^2^=737.31, P < 0.001 and for (b) χ^2^=810.53, P < 0.001)

How can the sequencing method have such an influence? It is important to highlight that high-throughput sequencing runs always produce spurious confidence reads that must be filtered out. In the case of short reads, this is done by imposing a minimum on the number of reads before calling an insertion site, whereas for long reads, this is done by imposing a minimum on the number of detected poly-A tails (see above). The fact that a poly-A tail is detected at all is already a token of quality, so an insertion site supported by one poly-A tail is inherently more robust than an insertion site supported by one read. Therefore, long-read technologies are favorable because they allow the real insertion sites to be identified more readily than with short reads.

Otherwise, the integration biases were reproducible among independent transductions with both Illumina (**Fig. 4b**) and Nanopore (**Fig. 4c**). This indicates that both sequencing technologies give similar large-scale results, even though the details of the insertion sites and their robustness differ. In conclusion, long-read technologies improve the robustness of Terminal mapping by leveraging a specific feature of the workflow. Short-read technologies impose a higher level of noise but they remain compatible with Terminal mapping.

### Terminal mapping identifies discrepancies between insertion profiles

Based on the benchmarking results above, we set Oxford Nanopore Technologies as the default sequencing platform and returned to the goal of establishing baseline insertion profiles for HIV-1 and SIVmac251 in Jurkat cells. The chromosomal distribution of integrated HIV-1 and SIVmac251 for all the independent transductions showed a clear enrichment on chromosomes 16, 17 and 19 (**Fig. 5a**), as observed by us and others (Chen et al., 2017; Marini et al., 2015; Vansant et al., 2020). The Circos plot makes it clear that chromosomes X and Y are targeted less frequently than autosomes. Jurkat T cells are male so they have only one X chromosome, accounting for the ∼2-fold difference of density with, say, chromosome 1 that is present in two copies. The density of insertions is much lower in the Y chromosome than in the X chromosome, most likely because the Y chromosome is unmappable due to the high content of repeated sequences. Interestingly, the insertion density on chromosome 18 is approximately the same as on chromosome X. It is known that one of the two chromosomes 18 is missing in Jurkat cells (Wilson et al., 2025). Therefore, DNA content seems to be a strong driver of the insertion signal, suggesting that the differences between Jurkat-a and Jurkat-b may be due to variations in their karyotypes — the karyotype of Jurkat cells is known to drift over time due to mutations in the DNA repair system and in checkpoint proteins (Gioia et al., 2018).

**Figure 5.**
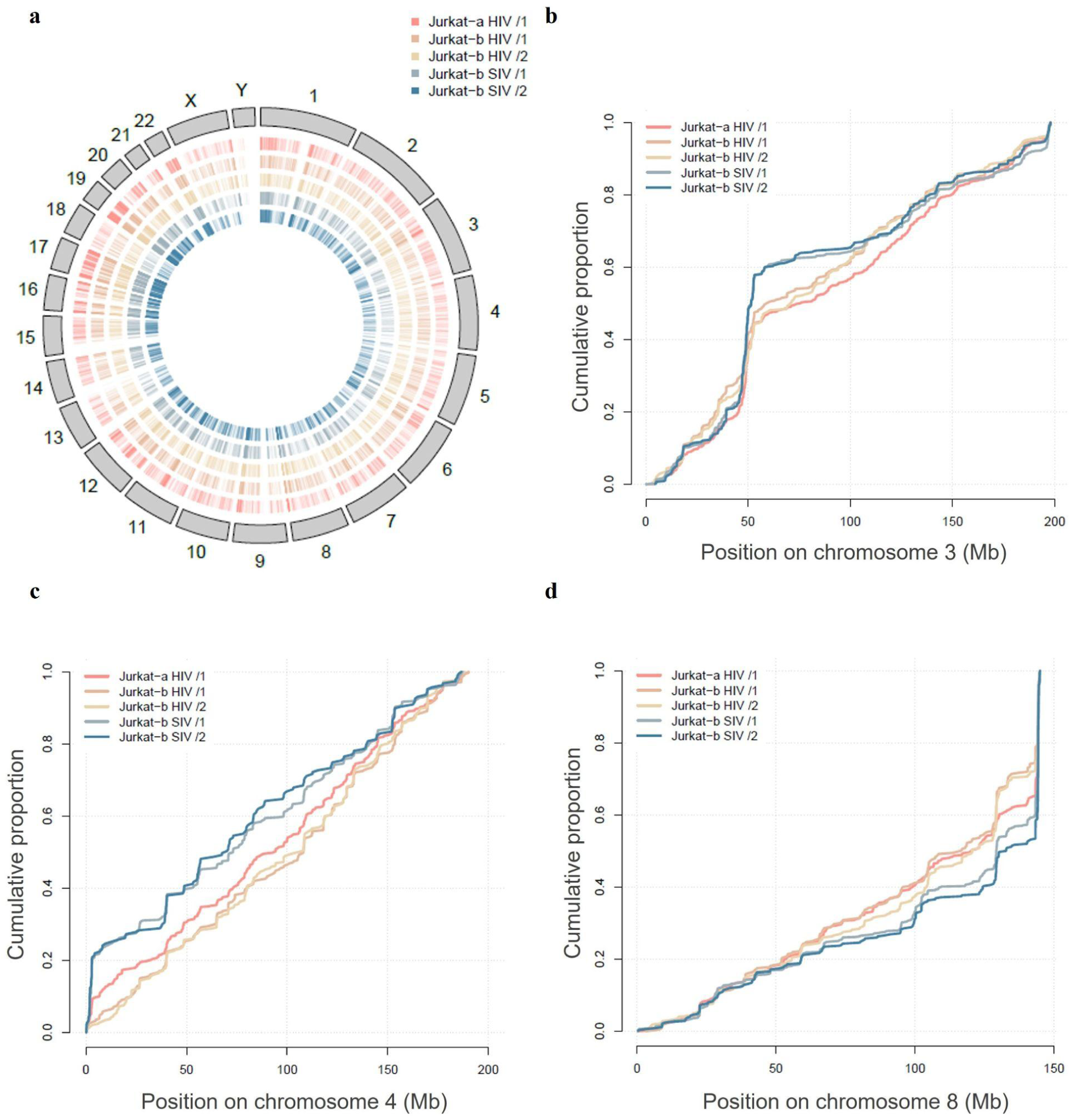
HIV-1 and SIVmac251 integration hotspots. (a) Genome-wide map of HIV-1 and SIVmac251 integrations in independent transductions using Jurkat T cells from different sources, Jurkat-a and Jurkat-b. The grey bars represent the human chromosomes, and the pink and blue ticks indicate the positions of HIV-1 and SIVmac251 pseudotyped virus integrations mapped by Terminal mapping in Jurkat-a and Jurkat-b, respectively. HIV-1 and SIVmac251 integration hotspots on chromosome 3 (b), chromosome 4 (c) and chromosome 8 (d) in Jurkat-a and Jurkat-b. For (b), (c) and (d), the lines show every integration with a jump of fixed height and hotspots appear as near-vertical lines. The differences between the HIV-1 and SIVmac251 constructs in Jurkat-b are all highly statistically significant (two-sided Kolmogorov-Smirnov test, chromosome 3: D=0.120 and P < 0.001, chromosome 4: D=0.198 and P < 0.001, chromosome 8: D=0.164 and P < 0.001). The differences between Jurkat-a and Jurkat-b for the HIV construct are statistically significant at level alpha=0.05 (two-sided Kolmogorov-Smirnov test, chromosome 3: D=0.080 and P=0.0108, chromosome 4: D=0.091 and P=0.0136, chromosome 8: D=0.099 and P=0.0024)

However, DNA content is not the sole driver of the signal, since HIV-1 and SIVmac251 have markedly different profiles in the same cell clone Jurkat-b (**Fig. 4b and 4c**). This shows that viruses also have markedly different preferences for certain regions. Most notable are chromosomes 16, 17 and 19, where the relative insertion rate is ∼50% higher for SIVmac251 than for HIV-1, which is on par with the differences between Jurkat-a and Jurkat-b. This indicates that insertion profiles at the scale of the genome are equally influenced by DNA content and by viral preferences.

To further study the integration preferences of HIV-1 vs SIVmac251, we plotted the cumulative distributions of the insertion sites on individual chromosomes (**Fig. 5b–d**). In this representation, the local slope of the curve indicates the insertion rate, so a near-vertical line indicates a hotspot and a horizontal line indicates an insertion desert. We observed visible differences on chromosome 3 (**Fig. 5b**), chromosome 4 (**Fig. 5c**) and chromosome 8 (**Fig. 5d**). Chromosomes 3 and 8 with 40% and 50% SIVmac251 integrations, respectively, show visible hotspots (**Fig. 5b and 5d**). The beginning of chromosome 4 (∼1.7% of the chromosome length), represents 20% of SIVmac251 integrations (the vertical line jumps to ∼20% on the y-axis) while HIV-1 barely has a hotspot at the same location in Jurkat-b and has a somewhat weaker hotspot in Jurkat-a (**Fig. 5c**). It is remarkable that SIVmac251 has a stronger hotspot on chromosomes 3 and 4, but it integrates less on both these chromosomes compared to HIV-1 (**Fig. 4b and 4c**), indicating that SIVmac251 is enriched in some places while being depleted in others. Otherwise, it is remarkable that the main visible differences between HIV-1 and SIVmac251 are in the strength of the hotspots but not their locations. A hotspot for HIV-1 is usually a hotspot for SIVmac251 and conversely, but to a different degree. In all the cases that we observed, hotspots were stronger for SIVmac251, indicating a bimodal insertion profile with hotspots and insertion deserts, but few regions of intermediate insertion rate.

Taken together, these results show that Terminal mapping can be used to identify differences of insertion profiles at the genome and at the chromosome levels. In addition, we have shown that insertion profiles can differ based on the cell clone, probably due to changes of DNA content, and they can differ based on the virus, due to insertion preferences. We showed that HIV-1 and SIVmac251 share the same insertion hotspots, but the profiles are distinct because the strength of the hotspots varies to a large extent.

### Terminal mapping can be used to monitor unintegrated LTR viruses

An interesting property of Terminal Mapping is that the primer for linear amplification anneals to both 5’ and 3’ LTRs (**Fig. 2**). This is necessary because the sequences of the LTRs are identical after the insertion into the host genome (barring mutations). In principle, Terminal Mapping can be performed on any kind of mobile DNA element, but in the case of LTR viruses, this property gives additional leverage for internal controls. We have seen above that it is possible to use the products from the 3’ LTR to assess the progress of the linear amplification, as in **Supplementary Fig. 2**. In addition, those products can also be used after sequencing to verify that both LTRs produce roughly the same amount of reads. This is expected to be the case at equilibrium when proviruses are stably integrated, but not when a large number of viruses are not integrated, *e.g.*, shortly after the infection. Indeed, unintegrated viruses including pre-integration viruses, one-LTR and two-LTR circles, produce reads from the 3’ LTR that can be mapped to the virus, while the reads from the 5’ LTR cannot be mapped to the host genome (because the 5’ LTR is not next to a sequence of the host in any of those cases). As a result, it is possible to compare the number of reads from the 5’ and 3’ LTRs to establish whether unintegrated viruses have been washed out.

To demonstrate this, we collected the proportions of reads from the 3’ LTR in the experiments presented above. We obtained 51% for HIV-1 in Jurkat-a, 46% and 45% for HIV-1 in Jurkat-b and 51% and 52% for SIVmac251 in Jurkat-b (Methods). These numbers show that there are approximately the same number of reads coming from both LTRs, suggesting that in the conditions of the experiment (*i.e.*, 21 days post transduction), unintegrated viruses have been completely washed out.

To test the methodology, we performed Terminal Mapping on Jurkat-b four days post transduction with pseudotyped reporter HIV-1 viruses at an MOI of 1.0, without FACS sorting GFP-positive cells. In these conditions, it is expected that unintegrated viruses are present in the cells and can even dominate (Suspène & Meyerhans, 2012). Over four independent infections, the proportion of reads from the 3’ LTR was found to be 96%, 93%, 98% and 97%. These results demonstrate that the proportion of reads from the 3’ LTR is indeed higher shortly after the infection, thereby confirming that this can be used as an internal control to ascertain that the viruses that can be amplified by PCR are integrated into the host genome.

Taken together, these results indicate that Terminal Mapping can be used to monitor the proportion of unintegrated viruses. This in turn is useful when measuring viral expression shortly after the infection, as it allows experimenters to estimate the breakdown of mRNA or protein originating from integrated vs unintegrated viruses.

More generally, our results collectively indicate that Terminal Mapping is a robust mapping technique that has several advantages over existing methods, including internal controls that can be used to further increase robustness with long reads or to monitor unintegrated viruses.

## DISCUSSION

Here, we developed a novel targeted sequencing approach called Terminal Mapping which combines linear amplification and poly-A-tailing followed by high-throughput sequencing to precisely identify proviral integration sites in Jurkat T cell lines transduced with HIV-1 or SIVmac251 based vectors. Libraries were sequenced using Illumina and Nanopore technologies and mapping workflows with short and long reads, respectively, were compared. We recapitulated the known features of HIV-1 integration that we had found in one of our previous works using the Jurkat T cell line-a (Chen et al., 2017). We also determined the first of its kind high-resolution hotspot map of SIVmac251 integration sites. Furthermore, we also show that the HIV-1 integration landscape has strikingly large differences between two batches of the Jurkat T cell line, Jurkat-a and Jurkat-b. Overall, these results illustrate the power of the novel high-throughput mapping method described in this study and highlight the importance of cell-to-cell variability in HIV-1 persistence.

Previous studies have shown that SIV and HIV-1 share integration target site preferences, implying similar mechanisms for selecting target sites by these lentiviruses (Crise et al., 2005; Ferris et al., 2019). The general consensus regarding HIV-1 and SIV pre-integration complexes (PICs) is that they interact with the same host factors as the PICs transit the cytoplasm, cross the nuclear membrane, and move through the nucleus to associate with the chromatin that contains the preferred integration sites (Lee et al., 2010). In our study, the overall distribution of HIV-1 and SIVmac251 integration sites was found to be quite similar with strong integration bias towards chromosomes 16, 17 and 19. Detailed observation shows differences in the integration hotspots of HIV-1 and SIVmac251 proviruses on chromosomes 3, 4 and 8. For example, at the beginning of chromosome 4 (∼1.7% of the chromosome length), SIVmac251 has 20% integration while HIV-1 barely has a hotspot at the same location in Jurkat-b and has a somewhat weaker hotspot in Jurkat-a.

The differences between the insertion profiles of HIV-1 constructs in Jurkat-a vs Jurkat-b are remarkable at the global scale. For instance, the insertion rate in chromosome 19 is ∼40% higher in Jurkat-a than in Jurkat-b. Prior to this study, we worked since 2013 with Jurkat-a, and in our experience the insertion profiles were always reproducible, even when transductions were performed several years apart (Chen et al., 2017; Vansant et al., 2020). Upon establishing our laboratory at the University of Toronto, we acquired a new source of Jurkat cells from the clone E6-1 (Jurkat-b) and immediately noticed the difference. Unfortunately, only one transduction performed in Jurkat-a remained in our DNA stocks, so it was impossible to obtain a second replicate of the insertion profile in the present study. However, this single replicate is similar to the ones we obtained while working with Jurkat-a.

Surprisingly, those large differences at the global level do not seem to emerge from different insertion profiles on the chromosomes. For the HIV-1 constructs, the cumulative insertion profiles on chromosomes 3, 4 and 8 follow similar trajectories, indicating that the patterns of enrichment and depletion are similar between Jurkat-a and Jurkat-b. The other chromosomes are not shown, but the results are similar (the largest differences were found on chromosomes 3, 4 and 8). The insertion hotspots at the leftmost part of chromosome 4 and at the rightmost part of chromosome 8 are stronger in Jurkat-a than in Jurkat-b (the cumulative profiles show a bigger vertical jump), but the overall insertion rate of HIV-1 on chromosomes 3 and 8 is actually lower in Jurkat-a than in Jurkat-b. Thus, it seems that chromosomes are uniformly heightened or lowered, meaning that the insertion rate in a chromosome can change between Jurkat-a and Jurkat-b, but the insertion profile remains similar.

This may suggest that Jurkat-a and Jurkat-b have different karyotypes, whereby Jurkat-a has extra chromosomes 16, 17 and 19 compared to Jurkat-b. This is particularly relevant as Jurkat clones have an aberrant karyotype and are prone to chromosomal instability. Jurkat cells were isolated in 1977 from a patient with T-cell acute lymphoblastic leukemia. Like for many cancer cells, the genome of Jurkat cells contains chromosomal rearrangements and mutations in maintainers of genome integrity such as *TP53* (p53) and *CDKN2A* (p16). Importantly, chromosomes 17 and 19 are often overrepresented and rearranged, explaining to some extent the skewness of the global insertion profile of HIV-1 constructs. Therefore, it is plausible that the differences of insertion rate between Jurkat-a and Jurkat-b coincide with copy-number variations. However, it is impossible to conclude definitively in the absence of more precise karyotypic data for Jurkat-a and Jurkat-b.

That said, it is important to mention that karyotypic differences cannot explain all the features of the insertion pattern. Focusing on the differences between the HIV-1 and SIVmac251 constructs, it appears that large variations are possible at the global level (*e.g*., the insertion rate on chromosome 19) or at the chromosomal level (*e.g*., the hotspots on chromosomes 3, 4 and 8). Those variations taking place solely within Jurkat-b are wider than the variations observed for HIV-1 constructs between Jurkat-a and Jurkat-b. This shows that the nature of the viral constructs and its interactions with the host cell can have bigger effects than differences of karyotype. Therefore, it is plausible that the observed differences between Jurkat-a and Jurkat-b are due in part to karyotypic differences, and in part to other factors changing the interactions of the HIV-1 construct and the host cell.

A major consideration is that our work is based on experiments performed on a single immortalized cell line, Jurkat T cell line. Jurkat T cell cultures are not homogeneous populations. Instead, they comprise multiple sub-clones that differ in transcriptional activity and proliferative capacity, which profoundly influences HIV-1 integration site selection and proviral expression (Skupsky et al., 2010; Atindaana et al., 2022). In a previous study, Janssens *et al*. have directly visualized heterogeneous subpopulations in Jurkat T cells using single-cell imaging, demonstrating that integration into active chromatin correlates with higher reactivation potential, whereas clones harboring proviruses in repressive regions remain largely latent (Janssens et al., 2022). The evidence that HIV-1 has evolved autonomous mechanisms to shut down its own expression (Razooky et al., 2015) does not exclude the role of the genomic environment emerging as a significant control parameter for gene expression variation (Skupsky et al., 2010). This is of particular importance because latent HIV-1 infections do not induce cellular responses, underscoring that the observed variability stems from pre-existing clonal differences rather than infection-driven transcriptional reprogramming (Matsui et al., 2021). The intrinsic heterogeneity of Jurkat T sub-clones highlights the magnitude of research that is necessary to investigate the distribution of HIV-1 integration sites and gene expression across the cell culture. At a larger scale, the heterogeneous population of Jurkat T cell line mimics the multiple heterogeneous tissue-sequestered and circulating subsets of CD4+ T cells, each with distinct epigenetic states and transcriptomic signatures.

The findings of this study must be seen in light of a few relevant limitations. The pseudoviruses used in this work were generated as single-round, VSV-G-pseudotyped particles produced by co-transfecting a minimal transfer plasmid together with a packaging construct that supplies the structural proteins and HIV-1 integrase in trans. Because the integrase enzyme is supplied from the packaging plasmid rather than being encoded in the viral genome, its expression level, timing and stoichiometry may differ from that of a native, full-length HIV-1 or SIV virion. This simplification can alter nuclear import dynamics and the chromatin contexts encountered by the pre-integration complex, potentially biasing the observed integration site profile. The insertion site profiles obtained in the cell-line model may not capture all virus-specific biases present in natural infection. Nevertheless, the core determinants of retroviral integration such as preference for transcriptionally active chromatin and association with specific epigenetic marks are preserved, allowing the data to inform how these genomic features guide proviral integration in primary cells and clinical samples.

Our results indicate that Terminal Mapping benefits from long reads, but a full discussion of sequencing technologies goes beyond read length and associated quality controls. A hallmark of the Illumina technology is that the sequencers with highest throughput require heavy infrastructure, to the effect that they are typically embedded in facilities such as sequencing centers. This in itself adds a layer of logistics, as samples must be queued, typically for weeks, before they can be sequenced. Some Illumina sequencers, such as the NextSeq series, have an increased turn-over due to their altered chemistry optimized for speed, but they are still mostly embedded in sequencing centers. Importantly, the modified chemistry of the NextSeq disallows sequencing PCR amplicons with constant sequence fragments (Shendure et al., 2019), such as the LTR or the PCR primers, that are present in all the products of Terminal Mapping. There are ways to bypass the limitations, but this requires setting up non-standard sequencing runs with special primers, which further increases the logistic burden. In summary, the high throughput of the Illumina technology comes at the cost of a long turn-around time that slows down troubleshooting and methodological development. For Illumina sequencing, libraries were sent to a sequencing center. Like with any other sequencing facility, waiting periods ranging from weeks to months to obtain the raw reads and high cost remained a challenge.

In comparison, the Oxford Nanopore technology is better suited for a method in development because sequencers are portable and fit on a bench top, samples do not need to be shipped and library preparation can be done in under one hour. Samples can still be queued for days, but the wait time is usually lower than for Illumina machines. The downside of the Oxford Nanopore technology is the higher error rates, making it challenging to map reads or to identify key features such as LTRs or other sequences embedded in the reads. An additional complication is that the library preparation for PCR amplicons requires the ligation of sequencing adapters. This step is somewhat imperfect and it tends to generate concatemers, whereby multiple PCR amplicons are ligated in tandem. This is not a problem, as the amplicons are all sequenced, even if the concatemer is large, but this adds to the difficulty of analyzing the results, as concatemers must be disambiguated. In conclusion, we opted for the Oxford Nanopore technology in order to shorten the turnaround time and to troubleshoot as fast as possible, but we had to develop bespoke computational pipelines to analyze and interpret the results.

We acknowledge that Terminal Mapping is not yet ready for the clinic, and the current prototype should only be used for scientific research. Firstly, the protocols were developed for viral constructs with a known sequence, whereas viruses in the wild can mutate and diverge to the point that they are not amplified by a reference primer for linear amplification. This is especially true of HIV or SIV that tend to mutate faster than other viruses. Secondly, we have indirect evidence that the current prototype lacks the sensitivity to pick up every individual insertion of the virus in the sample. The purpose of starting Terminal Mapping with 100 steps of linear amplification is to boost the signal so that every single provirus molecule gives ∼100 DNA copies that can be mapped independently. However, in the results, we typically find 1–3 DNA copies per provirus. This can be inferred from the number of distinct positions where a poly-A tail is found in the sequencing results. This shows that most DNA copies are lost or not created at all during the Terminal Mapping process, suggesting that many insertion sites may be missing if they are present as a single copy in the sample. Overall, it is likely that the purification steps of Terminal Mapping must be further optimized if it were to pick up every single insertion site. Finally, Terminal Mapping is a cost-effective protocol for preparing libraries that has the potential to characterize highly complex samples with multiple target sequences and has sufficient sensitivity, specificity, and robustness to detect and sequence the relatively rare integration sites via an initial linear amplification step, highlighting an application (with further optimization) on clinical samples. Due to high sensitivity, Terminal Mapping is prone to contamination if executed inattentively. Thus, a PCR-grade environment and special attention to clean handling of the protocol is of utmost importance to successfully amplify the unknown flanking DNA without contaminating samples. Therefore, including negative controls into the protocol is strongly recommended. If control samples indicate that cross-contamination occurred during the protocol, the products from every pause point can be used to repeat parts of the protocol. Our goal with this approach is to allow library preparation without requiring any DNA fragmentation, which can waste large amounts of input material (when using sonication) or limit the range of mappable insertions (when using restriction enzymes), a step replaced by the generation of poly-A-tail using the enzyme Terminal transferase. We foresee the application of Terminal Mapping, with some minor modifications, to map any unknown DNA sequences adjacent to a known DNA sequence, offering a relatively rapid ready-to-use high-throughput technology.

In summary, we have developed a novel technology called Terminal Mapping to map mobile elements and have tested it in Jurkat cells on LTR constructs derived from HIV-1 and SIVmac251. We showed that Terminal Mapping is both sensitive and robust, and that its workflow leverages long-read technologies. Using Terminal Mapping, we showed that HIV-1 and SIVmac251 share similar hotspots but have distinct insertion profiles, demonstrating that the method can be applied to compare profiles, with promising applications for biotechnologies related to mobile elements.

## SUPPLEMENTARY MATERIALS

**Supplementary Figure 1.**
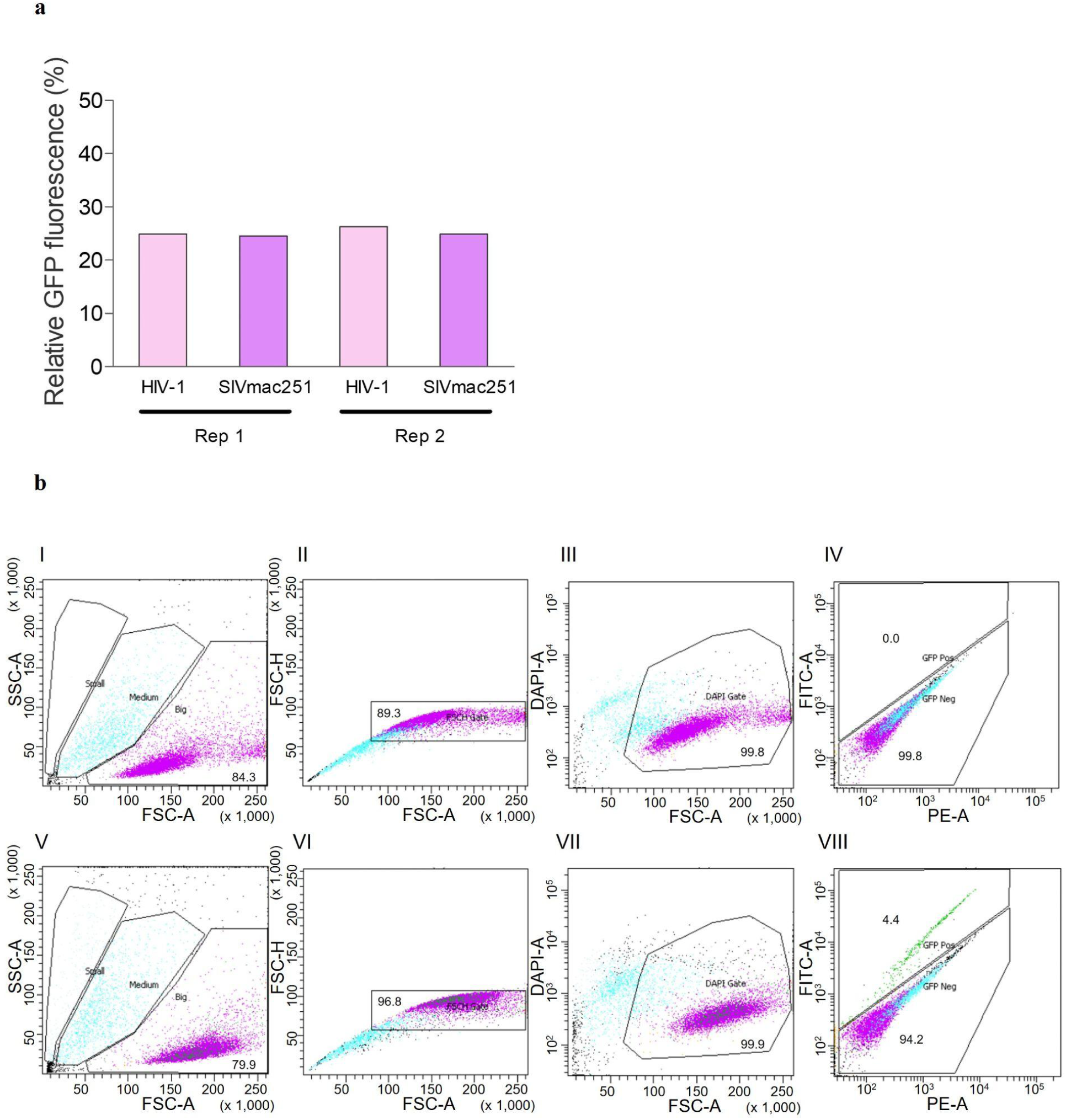
Efficiency of HIV-1 and SIVmac251 transduction in Jurkat T cells and FACS isolation of GFP(+) transduced cells. (a) Transduction efficiency of minimal HIV-1 and SIVmac251 construct in Jurkat T cells were checked by measuring GFP expression 48 hours post transduction by FACS analysis. Experiments were performed twice. (b) Representative FACS profiles for sorting GFP (+) cells 48 hours post HIV-1 or SIVmac251 transduction (bottom panel) and untransduced negative control cells (top panel). (I and V) Jurkat cells were selected by FSC-A versus SSC-A (Big gated region). (II and VI) Singlets were selected by FSC-A versus FSC-H (gated region). (III and VII) Live cells (gated region) were selected by DAPI (-) on FSC-A versus DAPI-A. (IV and VIII) GFP (+) cells were acquired by GFP fluorescence. Negative control did not have GFP (+) cells showing successful transduction.

**Supplementary Figure 2.**
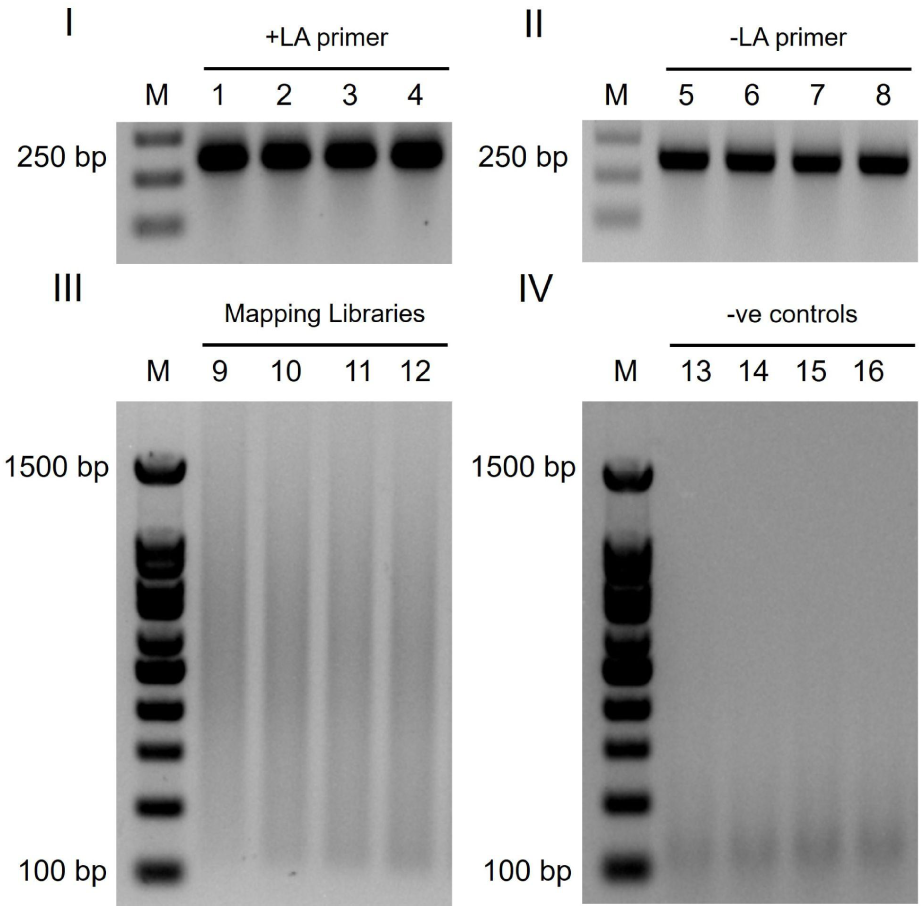
Validation of Terminal Mapping. Representative PCR products were loaded on 2.0% (wt/vol) agarose gel. I – The expected size of PCR amplicons to confirm linear amplification was 250 bp. Lane 1,2,3 and 4: Linear amplified product; II - Lane 5,6,7 and 8: No linear amplification control. III – PCR smears were expected to lie between 100 bp and 1500 bp. The PCR smear corresponds to different integrations in Jurkat T cells either transduced by HIV-1 or SIVmac251 (lane 9,10,11 and 12). The smears were specific, because they failed to appear when the cells were not transduced (lane 13) or when linear amplification was not performed (lane 14) or when poly-A-tailing was not performed (lane 15) or when templates were not added (lane 16).

**Supplementary Figure 3.**
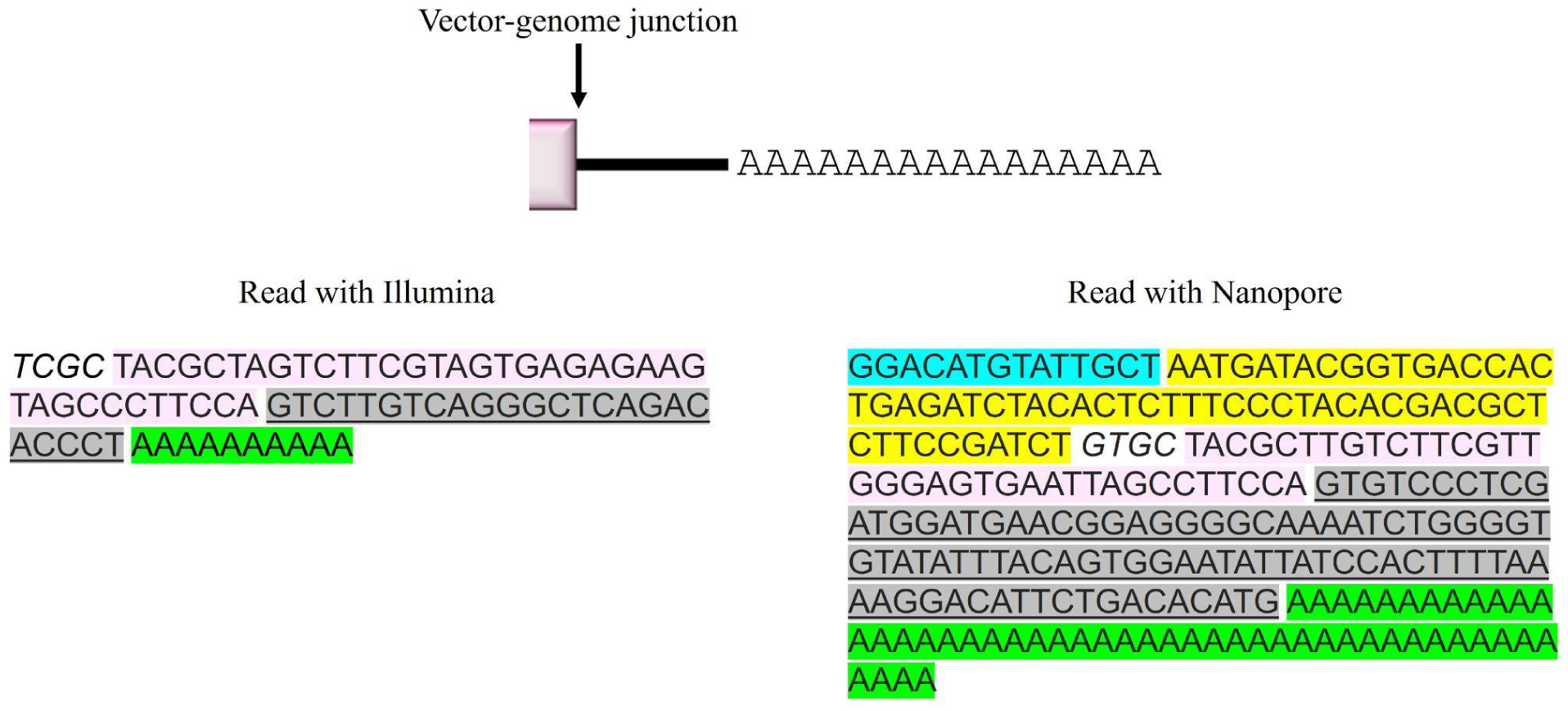
Comparison of the HIV-1 integration site analysis between Illumina and Oxford Nanopore sequencing technology. Representation of a typical HIV-1 read pattern obtained from Illumina and Oxford Nanopore sequencing technology. The reads obtained from Illumina are shorter compared to the ones obtained from Nanopore. The difference in the read length between the two platforms plays a significant role for the downstream analysis of the results. Left - A typical read with Illumina. First 4 random nucleotides; Nucleotides with pink background represent sequence from a HIV-1 vector; Nucleotides with grey background represent sequence from a putative human genome part; Nucleotides with green background represent sequence from a poly-A-tail added by the terminal transferase enzyme. Right - A typical read with Nanopore. Nucleotides with blue background represent sequence from the Nanopore adapter; Nucleotides with yellow background represent sequence from the PCR adapter; 4 random nucleotides; Nucleotides with pink background represent sequence from a HIV-1 vector; Nucleotides with grey background represent sequence from a putative human genome part; Nucleotides with green background represent sequence from a poly-A-tail added by the terminal transferase enzyme. Pink box: Partial of HIV-1 5’LTR; black line: human genome.

---

**Supplementary Table 1.**
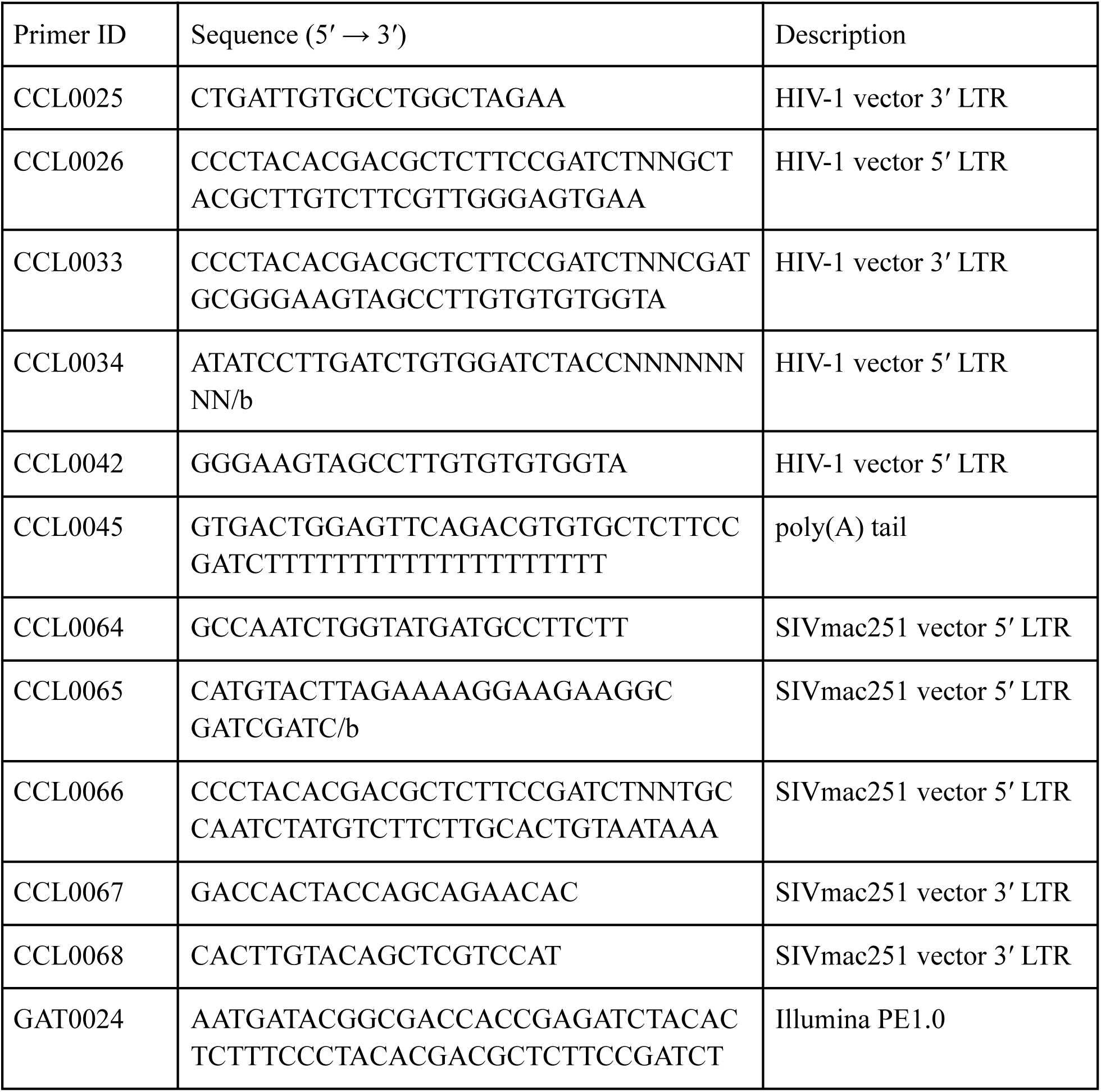
List of primers used in Terminal Mapping.

**Supplementary Table 2.**
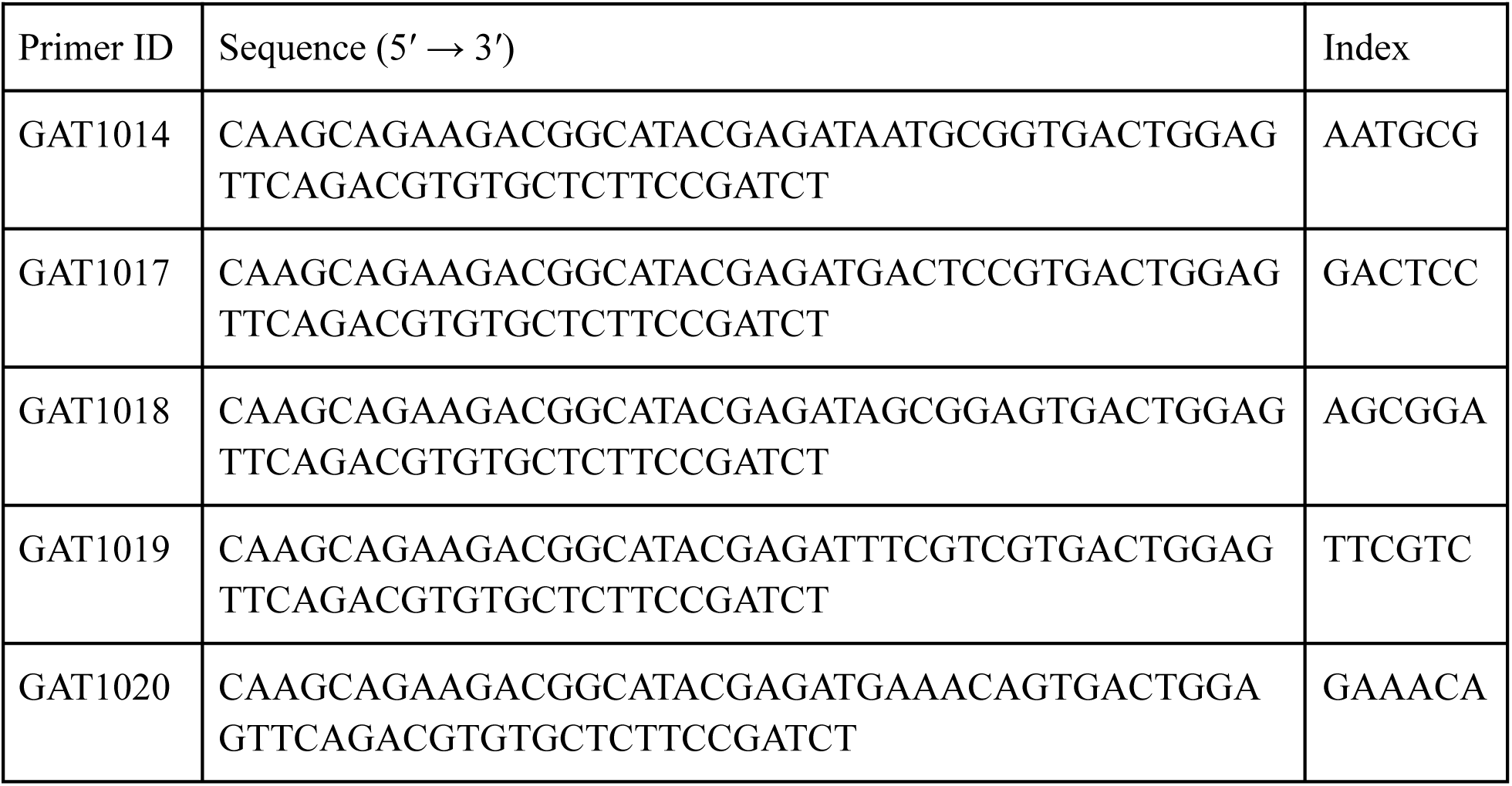
List of indexing primers used in Terminal Mapping.

## REFERENCES

Atindaana, E., Kissi-Twum, A., Emery, S., Burnett, C., Pitcher, J., Visser, M., Kidd, J. M., & Telesnitsky, A. (2022). Bimodal Expression Patterns, and Not Viral Burst Sizes, Predict the Effects of Vpr on HIV-1 Proviral Populations in Jurkat Cells. mBio, 13(2), e0374821.

Bulcha, J. T., Wang, Y., Ma, H., Tai, P. W. L., & Gao, G. (2021). Viral vector platforms within the gene therapy landscape. Signal Transduction and Targeted Therapy, 6(1), 53.

Bushman, F., Lewinski, M., Ciuffi, A., Barr, S., Leipzig, J., Hannenhalli, S., & Hoffmann, C. (2005). Genome-wide analysis of retroviral DNA integration. Nature Reviews Microbiology, 3(11), 848–858.

Chen, H.-C., Martinez, J. P., Zorita, E., Meyerhans, A., & Filion, G. J. (2017). Position effects influence HIV latency reversal. Nature Structural & Molecular Biology, 24(1), 47–54.

Crise, B., Li, Y., Yuan, C., Morcock, D. R., Whitby, D., Munroe, D. J., Arthur, L. O., & Wu, X. (2005). Simian immunodeficiency virus integration preference is similar to that of human immunodeficiency virus type 1. Journal of Virology, 79(19), 12199–12204.

Ferris, A. L., Wells, D. W., Guo, S., Del Prete, G. Q., Swanstrom, A. E., Coffin, J. M., Wu, X., Lifson, J. D., & Hughes, S. H. (2019). Clonal expansion of SIV-infected cells in macaques on antiretroviral therapy is similar to that of HIV-infected cells in humans. PLoS Pathogens, 15(7), e1007869.

Gioia, L., Siddique, A., Head, S. R., Salomon, D. R., & Su, A. I. (2018). A genome-wide survey of mutations in the Jurkat cell line. BMC Genomics, 19(1), 334.

Goujon, C., Jarrosson-Wuillème, L., Bernaud, J., Rigal, D., Darlix, J.-L., & Cimarelli, A. (2006). With a little help from a friend: increasing HIV transduction of monocyte-derived dendritic cells with virion-like particles of SIV(MAC). Gene Therapy, 13(12), 991–994.

Hickey, D. A. (1982). Selfish DNA: a sexually-transmitted nuclear parasite. Genetics, 101(3-4), 519–531.

Janssens, J., De Wit, F., Parveen, N., & Debyser, Z. (2022). Single-Cell Imaging Shows That the Transcriptional State of the HIV-1 Provirus and Its Reactivation Potential Depend on the Integration Site. mBio, 13(4), e0000722.

Jordan, M., Schallhorn, A., & Wurm, F. M. (1996). Transfecting mammalian cells: optimization of critical parameters affecting calcium-phosphate precipitate formation. Nucleic Acids Research, 24(4), 596–601.

Lee, K., Ambrose, Z., Martin, T. D., Oztop, I., Mulky, A., Julias, J. G., Vandegraaff, N., Baumann, J. G., Wang, R., Yuen, W., Takemura, T., Shelton, K., Taniuchi, I., Li, Y., Sodroski, J., Littman, D. R., Coffin, J. M., Hughes, S. H., Unutmaz, D.,…KewalRamani, V. N. (2010). Flexible use of nuclear import pathways by HIV-1. Cell Host & Microbe, 7(3), 221–233.

Lusic, M., & Siliciano, R. F. (2017). Nuclear landscape of HIV-1 infection and integration. Nature Reviews. Microbiology, 15(2), 69–82.

Madhani, H. D. (2013). The frustrated gene: origins of eukaryotic gene expression. Cell, 155(4), 744–749.

Mangeot, P.-E., Duperrier, K., Nègre, D., Boson, B., Rigal, D., Cosset, F.-L., & Darlix, J.-L. (2002). High levels of transduction of human dendritic cells with optimized SIV vectors. Molecular Therapy: The Journal of the American Society of Gene Therapy, 5(3), 283–290.

Marini, B., Kertesz-Farkas, A., Ali, H., Lucic, B., Lisek, K., Manganaro, L., Pongor, S., Luzzati, R., Recchia, A., Mavilio, F., Giacca, M., & Lusic, M. (2015). Nuclear architecture dictates HIV-1 integration site selection. Nature, 521(7551), 227–231.

Matsui, H., Shirakawa, K., Konishi, Y., Hirabayashi, S., Sarca, A. D., Fukuda, H., Nomura, R., Stanford, E., Horisawa, Y., Kazuma, Y., Matsumoto, T., Yamazaki, H., Murakawa, Y., Battivelli, E., Verdin, E., Koyanagi, Y., & Takaori-Kondo, A. (2021). CAGE-seq reveals that HIV-1 latent infection does not trigger unique cellular responses in a Jurkat T cell model. Journal of Virology, 95(8). 10.1128/JVI.02394-20

Mueller, P. R., & Wold, B. (1989). In vivo footprinting of a muscle specific enhancer by ligation mediated PCR. *Science (New York*, N.Y.), 246(4931), 780–786.

Ochman, H., Gerber, A. S., & Hartl, D. L. (1988). Genetic applications of an inverse polymerase chain reaction. Genetics, 120(3), 621–623.

Razooky, B. S., Pai, A., Aull, K., Rouzine, I. M., & Weinberger, L. S. (2015). A hardwired HIV latency program. Cell, 160(5), 990–1001.

Schmidt, M., Schwarzwaelder, K., Bartholomae, C., Zaoui, K., Ball, C., Pilz, I., Braun, S., Glimm, H., & von Kalle, C. (2007). High-resolution insertion-site analysis by linear amplification-mediated PCR (LAM-PCR). Nature Methods, 4(12), 1051–1057.

Schröder, A. R. W., Shinn, P., Chen, H., Berry, C., Ecker, J. R., & Bushman, F. (2002). HIV-1 integration in the human genome favors active genes and local hotspots. Cell, 110(4), 521–529.

Shendure, J., Balasubramanian, S., Church, G. M., Gilbert, W., Rogers, J., Schloss, J. A., & Waterston, R. H. (2019). Publisher Correction: DNA sequencing at 40: past, present and future. Nature, 568(7752), E11.

Skupsky, R., Burnett, J. C., Foley, J. E., Schaffer, D. V., & Arkin, A. P. (2010). HIV promoter integration site primarily modulates transcriptional burst size rather than frequency. PLoS Computational Biology, 6(9). 10.1371/journal.pcbi.1000952

Suspène, R., & Meyerhans, A. (2012). Quantification of unintegrated HIV-1 DNA at the single cell level in vivo. PloS One, 7(5), e36246.

Triglia, T., Peterson, M. G., & Kemp, D. J. (1988). A procedure for in vitro amplification of DNA segments that lie outside the boundaries of known sequences. Nucleic Acids Research, 16(16), 8186.

Vansant, G., Chen, H.-C., Zorita, E., Trejbalová, K., Miklík, D., Filion, G., & Debyser, Z. (2020). The chromatin landscape at the HIV-1 provirus integration site determines viral expression. Nucleic Acids Research, 48(14), 7801–7817.

Wilson, A., Dasyam, N., O’Hagan, F., Farrell, D., Turner, C., Jay, A., Schmidt, A., Beddow, R., Lillis, T., Chin, T., Perret, R., McCulley, M., & Weinkove, R. (2025). Jurkat T-cell lines exhibit marked genomic instability affecting karyotype, mutational profile, gene expression, immunophenotype and function. Scientific Reports, 15(1), 22426.

